# Dissection of function and recognition mechanism of *M. tuberculosis* ESX-1 secreted virulence factor EspC

**DOI:** 10.1101/2021.09.24.461649

**Authors:** Ruby Sharma, Vipin K. Kashyap, Manoj Kumar, Abhisheka Bansal, Ajay K. Saxena

## Abstract

*Mycobacterium tuberculosis* uses the ESAT-6 system-1/type VII (ESX-1) system for secretion of virulence proteins into the host cell, however the mechanism of virulence proteins secretion, molecular components and regulation of ESX-1 system are only partly understood. In the current study, we have analyzed the biological function and recognition mechanism between ESX-1 virulence EspC and EccA_1_ ATPase proteins. The EspC enters into A549 human lung carcinoma cells and exhibited cytotoxicity, as observed in MTT Assay. To understand the recognition mechanism between EspC and EccA_1_ ATPase, the EspC and EccA_1_ mutants were generated based on EspC~EccA_1_ interactions, as observed in molecular modeling. Binding analysis shows that EspC export arm interacts specifically to the β-hairpin insertion motif of the TPR domain of EccA_1_ ATPase. Mutations in these epitopes lead to significant decrease/or abolish the binding between EspC and EccA_1_ ATPase. Our study provides insight into biological function and recognition mechanism between EspC and EccA_1_ ATPase, which can be used as target to prevent EspC secretion/ or in general virulence factor secretion by mycobacterial ESX-1 system.

---

*Mycobacterium tuberculosis* ESX-1 secretion system is involved in essential pathogenesis, *e.g*., host phagosomal permeabilization, mycobacterial escape to cytosol and macrophage killing [1–5]. The ESX-1 secretion system exports the virulence proteins across the lipid-rich cell wall and helps in the permeabilization of host macrophage phagosomal membrane. The ESX-1 virulence proteins are involved in pore-formation in the membrane, modulation of extracellular signaling pathway [6–10] and act as a structural component of extracellular region of secretion apparatus itself [11]. The RD1 (region of difference) genes and its surrounding regions *e.g*., extended RD1 [12, 13] in *M. tuberculosis* and *M. marinum* encode virulence proteins of the ESX-1 secretion system. The EspC (*Rv3615c*) gene is encoded in the extended RD1 locus and is also present in *Bacillus Calmette–Guérin* vaccine [14]. The EspC is highly immunodominant like ESAT-6 and CFP-10 proteins in latent and active tuberculosis infections [15]. The EspC is upregulated at low pH [16] and in low iron conditions [17], as observed *in vitro* transcriptional response study. In a recent study [18], the EspC is found co-precipitate with EspA in both cytosolic and membrane fraction of *M. tuberculosis* cells. The EspC exists as cluster or long filament [18] and C-terminal region of EspC is involved in the polymer formation, which causes the mycobacterial virulence.

Mechanism of EspC secretion and its involvement in the regulation of ESX-1 system is currently unknown. The ESX-1 virulence proteins contain general “secretion signal”, which targets the AAA^+^ ATPase before exporting across the mycobacterial inner membranes [19, 20]. The structure of “secretion signal” is identified in the crystal structures of *M. tuberculosis* ESAT-6/CFP-10 complex [21–25] and EspB protein [26]. The nuclear magnetic resonance [27], X-ray crystallographic [21, 22, 24] and yeast two-hybrid [19] studies have shown that export arm of virulence proteins are stabilized by interaction with AAA^+^ATPase [30], a key phase in the export cycle and a potential determinant of substrate specificity. Recently, the crystal structure of *Thermonospora curvata* EccC enzyme in complex with signal sequence of *Thermonospora curvata* EsxB had been determined [28]. In this structure, the C-terminal signal sequence of *T. curvata* EsxB (96-103 residues) forms a short amphiphatic helix that interacts with hydrophobic pocket of ATPase_3_ domain of *T. curvata* EccC enzyme. However, YxxxD secretion motif of EsxB signal peptide was found disordered in the crystal structure and appeared not involved in EccC recognition [28].

The EspC recognizes the EccA_1_ ATPase (*Rv3868*, 573 residues, Mw~67kD), an essential component of ESX-1 secretion system and is involved in mycobacterial survival and virulence proteins secretion [29]. The EccA_1_ ATPase mutants bind several enzymes involved in cell wall biogenesis and prevent mycolic acid synthesis [29]. The EccA_1_ ATPase contains N-terminal tetratricopeptide repeat (TPR) domain involved in ESX-1 virulence proteins recognition and the C-terminal ATPase domain involved in ATP hydrolysis [30]. The crystal structure of TPR domain of EccA_1_ has been determined (PDB-4F3V), which adopts a TPR like fold as observed in other TPR domain proteins [31]. Swapping of C-terminal region of EspC with CFP-10 impairs the EspC secretion by ESX-1 system, as EccC ATPase recognizes only the CFP-10 export arm [30].

Yeast two hybrid and genetic studies have been performed to understand the ESX-1 virulence proteins recognition by their targets. In a recent structural study on *T. curvata* EccC enzyme in complex with signal sequence of *T. curvata* EsxB [28] showed that signal sequence of *T. curvata* EsxB forms short amphipathic helix and binds to hydrophobic pocket observed at C-terminal of ATPase_3_ domain of EccC enzyme, which resulted in enhanced ATPase activity. Molecular basis of EspC recognition by EccA_1_ ATPase is currently unknown. It will be highly important to understand, how EspC export arm recognizes the TPR domain of EccA_1_ ATPase and interacting residues involved in binding between EspC and EccA_1_, as observed by Rosenberg *et. al*. [28].

In current study, we have purified and performed functional studies on *M. tuberculosis* EspC and EccA_1_ ATPase. To understand the mechanism of EspC~EccA_1_ recognition, we built the EspC~EccA_1_ complex model and interactions involved in EspC~EccA_1_ recognition were identified. The EspC and EccA_1_ mutants were generated based on these interactions and binding analysis was performed using wild type and mutant EspC and EccA_1_ proteins. CD spectroscopy was performed to analyze the conformation and stability of the wild type and mutant EspC and EccA_1_ proteins. Our study shows the biological activities and molecular mechanism involved in the EspC~EccA_1_ recognition, which is essential for EspC secretion by the ESX-1 secretion system.

## MATERIALS AND METHODS

### Chemicals and Reagents

All the reagents, of analytical grade were procured from Sigma-Aldrich.

### Preparation of wild type and mutant EspC proteins

The EspC gene encoding Met1-Thr103 residues was amplified from *M. tuberculosis H37Rv* strain and cloned into *pET21a*(+) vector. The EspC plasmid was transformed in *E. coli. BL21(DE3)* cells and culture was grown at 37°C in Luria Bertani media supplemented with 100 μg/ml ampicillin. The culture was grown till OD_600_ ~ 0.6-0.7 and induced with 1mM IPTG at 37°C. After induction, cells were grown for another 4 h and harvested by centrifugation at 16,000xg. The cell pellet was suspended in lysis buffer containing (20mM HEPES pH 7.5, 50mM NaCl, 5% glycerol, 4mM β-mercaptoethanol, 1mM phenylmethylsulfonylfluoride, 0.01%(w/v) Tween-20 and 0.01% (w/v) SDS) and disrupted by sonication. The disrupted lysate was centrifuged at 25000x g and supernatant was collected.

The supernatant was mixed with pre-equilibrated Ni-NTA resin (Sigma) and incubated for 1 h on 360° rocker at 4°C. The protein was loaded in an empty column and washed with buffer containing (20mM HEPES pH 7.5, 50mM NaCl, 5% Glycerol, 4mM β-mercaptoethanol, 1mM Phenylmethylsulfonyl fluoride, 0.01%(w/v) Tween-20, 0.01%(w/v) SDS and 10mM imidazole). The EspC was eluted in a buffer containing (20mM HEPES pH 7.5, 50mM NaCl, 5% Glycerol, 3mM β-mercaptoethanol, 1mM Phenylmethylsulfonyl fluoride, 0.01%(w/v) Tween-20, 0.01%(w/v) SDS, 100mM imidazole). The eluted fractions of EspC were concentrated and loaded on Superdex™200 HiLoad (16/60) column (GE Healthcare). The column was pre-equilibrated with buffer containing (20mM HEPES pH 7.5, 50mM NaCl, 5mM β-mercaptoethanol, 0.01%(w/v) Tween-20, 0.01%(w/v) SDS]. The peak fractions from size exclusion column were pooled and concentrated using 3 kD cutoff ultracentrifugal device (Millipore, USA). The EspC purity was checked on 12% SDS-PAGE and by mass spectrometry. The EspC concentration was measured using absorbance at 280 nm. The purified recombinant EspC contains 112 residues, Mw~11.9kD (extra Met residue at N-terminal, 103 residues of EspC and LEHHHHHH residues at C-terminal).

Two point mutants of EspC (EspC-D91A and EspC-Y89A) were generated using site directed mutagenesis technique and all mutants were confirmed by DNA sequencing. The EspC-Δ5, EspC-Δ10 and EspC-Δ20 truncated mutants were obtained using PCR based mutagenesis protocol and confirmed by gene sequencing. The wild type EspC plasmid was used as template for mutant PCR amplification. The amplified plasmid was incubated with DpnI enzyme for 1 h at 37°C and mutant plasmids were isolated using DH5α cell. All five EspC mutants were purified using the same protocol used for wild type EspC.

### MTT assay to examine the EspC cytotoxicity on A549 cells

The MTT assay was performed to evaluate the EspC cytotoxicity in human lung carcinoma A549 cells. The MTT assay detects the reduction of MTT [3-(4,5-dimethylthiazolyl)-2,5-diphenyl-tetrazolium bromide) by mitochondrial dehydrogenases, which forms the formazan product. The reaction gives the blue color, which measures the cytotoxicity or cell viability. The A549 cells were seeded in 96-well plates at a density of 15×10^4^ cells/well for 24 h. Afterwards, these cells were treated with EspC for various time points *e.g*., 2 h, 4 h, 6 h, 12 h and 24 h. After the treatment, 100 μl of MTT (total=2.5 mg) was added in each well containing A549 cells and was incubated at 37°C for 4 h. After incubation, the media was carefully aspirated from the wells and formazan crystals obtained in the wells were dissolved in 100 μl of DMSO. The solution was kept on shaker for 15 min to dissolve the entire crystals. All steps were performed under proper protection from light. The microplate reader (ThermoFisher Scientific multiplate reader) was used to measure the absorbance at wavelength 570 nm. The live cells were counted under an inverted phase contrast microscope in hematocytometer with 10X magnification.

### Confocal microscopy analysis

For confocal microscopy, the A549 cells were grown on coverslip and were washed three times with PBS buffer. Freshly prepared fixing solution (1% acetone in chilled Methanol) was added and incubated for 20 min at 4°C. The A549 cells were washed three times with PBS buffer for 3 min. The cells were permeabilized in PBS buffer (containing in 0.1% Triton X-100) for 5 min at room temperature. The cells were further washed three times with PBS buffer and blocked with 5% skimmed milk prepared in PBST buffer for 2 h. The cells were incubated with anti-His antibody (1:200 dilution) for 1 h followed by washing three times with PBS buffer for 5 min. FITC conjugated mouse secondary antibody (GeNei Laboratories, India) was then added (1:1000 dilution) and incubated for 1 h. The cells were again washed three times with PBS buffer for 5 min. 1 □l of DAPI (1μg/ml in PBST buffer) solution was added in the well and incubated for 10 min. The A549 cells were washed three times with PBST buffer for 5 min. Slides were labeled and a small drop of mounting agent (50% Glycerol in 1xPBS buffer) was added to it. Coverslip containing cells were kept upside down on mounting agent over the slide. The coverslip was then sealed with the nail paint. The slides were examined under confocal microscopy. Stained cells were imaged using Nikon confocal microscope at Central Instrumentation Facility (CIF) of Jawaharlal Nehru University.

### Western Blot analysis

To test the amount of EspC protein present in the A549 cells, we have performed the Western blot analysis. The A549 cells were treated with EspC protein (conc. ~ 100 μg/ml) for 2, 4, and 6 h. The treated cells were placed on ice and then washed with ice-cold PBS buffer. 70 μl of ice-cold lysis buffer (50mM Tris-HCl, pH 7.4, 150mM NaCl, 1mM EDTA, 1% Triton-X100, 0.1% SDS, 1mM PMSF, 1X proteinase inhibitor cocktail, 10% Glycerol) was added in it, followed by centrifugation at 12,000 x g for 20 min at 4°C. The supernatant was used for protein analysis by western blotting. The total amount of protein was estimated by Bradford assay. Total 80 μg protein was used for Western blot analysis.

The proteins were separated on SDS-PAGE and transferred on a PVDF membrane. The membrane was blocked for 1 h at room temperature using the 5% skimmed milk in TBST buffer. The membrane was incubated with anti-His antibody (1:2000 dilution) in 2.5% BSA in TBST buffer for overnight at 4°C. The membrane was washed three times with TBST buffer, 5 min each followed by incubation with HRP conjugated mouse secondary antibody (GeNei Laboratories, 1:2000 dilution) in 5% skimmed milk in TBST buffer at room temperature for 1 h. The membrane was further washed three times with TBST buffer for 5 min each. The membrane was developed using Luminata Forte (Merck Millipore, USA) followed by exposure on X-ray film.

### Preparation of wild type and mutant EccA_1_ proteins

The EccA_1_ gene encoding (Met1-Glu573) was amplified using *M. tuberculosis H37Rv* strain and cloned into *pET23a* (+) expression vector. The EccA_1_ plasmid was transformed into *E. coli BL21(DE3)* cells and culture was grown at 37°C in Luria Bertani media supplemented with 100 μg/ml ampicillin at 37°C, till OD_600_ reached to 0.6-0.7. The culture was induced with 0.5 mM IPTG at 22°C and grown further for another 10 h. The EccA_1_ was overexpressed in soluble fraction of cell. The cells were harvested by centrifugation at 16,000 x g and suspended in lysis buffer containing (20mM HEPES pH 7.5, 150mM NaCl, 5% Glycerol, 4mM β-mercaptoethanol, 1mM Phenylmethylsulfonyl fluoride, 4mM Benzamidine-hexachloride, 0.2mM MgCl_2_, 0.2mM ATP, 10mM Arginine and 0.2mg/ml Lysozyme). The cell lysate was centrifuged at 25,000 x g and supernatant was collected.

The supernatant was mixed with pre-equilibrated Ni-NTA resin (Sigma) and incubated 1 h at 360° rocker at 4 °C. The Ni-NTA column was washed with buffer containing (20mM HEPES pH 7.5, 150mM NaCl, 5% Glycerol, 4mM β-mercaptoethanol, 1mM Phenylmethylsulfonylfluoride, 3mM Benzamidinehexachloride, 10mM Arginine, 0.2mM MgCl_2_, 0.2mM ATP and 40mM imidazole). The EccA_1_ was eluted in buffer containing (20mM HEPES pH 7.5, 150mM NaCl, 5% glycerol, 4mM β-mercaptoethanol, 1mM Phenylmethylsulfonyl fluoride, 3mM Benzamidine hydrochloride, 10mM Arginine, 0.2mM MgCl_2_, 0.2mM ATP, and 250mM imidazole). The Ni-NTA eluted fractions were pooled, concentrated and loaded on Superdex™200 HiLoad (16/60) column (GE Healthcare). The column was pre-equilibrated with buffer containing (20mM HEPES pH 7.5, 150mM NaCl, 5% Glycerol, 1mM Phenylmethylsulfonyl fluoride, 10mM Arginine, 0.2mM MgCl_2_, 0.2mM ATP, 4mM β-mercaptoethanol). The peak fractions were pooled and concentrated using a 30kD cutoff ultracentrifugal device (Millipore, USA). The purity of EccA_1_ was analyzed on 12% SDS-PAGE and mass spectrometry. The concentration of protein was measured by absorbance at 280nm.

Two EccA_1_ mutants, W94A and Y87A were obtained using site directed mutagenesis technique. Wild type EccA_1_ plasmid encoding (Met1-Glu573 residues) was used as template for PCR amplification. The amplified EccA_1_ plasmid was digested with DpnI enzyme for 1 h at 37 °C. The digested EccA_1_ plasmid was transformed in DH5α□cells and both EccA_1_ mutants W94A and Y87A plasmids were isolated. The EccA_1_ mutant proteins were purified using the same protocol used for wild type EccA_1_ purification.

### EccA_1_ ATPase activity analysis

To analyze the ATPase activity, the EccA_1_ was dissolved in ATPase buffer containing 20mM HEPES pH 7.5, 5mM MgCl_2_. 1μl of (γ^32P^)ATP (0.1 mC_i_) was added in 10 ml of ATPase buffer containing 1.1 □g of EccA_1_ and incubated for different time intervals at 37°C. 1 μl of the reaction buffer was spotted on TLC plate after every 10 min. The TLC plate was developed in 0.5 M formic acid and 0.5 M LiCl and dried at 37 °C. The TLC plate was exposed to Fuji film BAS-MS 2025 imaging plate for 12 h and analyzed using Typhoon FLA-9500 (from GE Healthcare). The background obtained from reaction buffer (having no EccA_1_ and (γ^32P^) ATP) was corrected in each ATP hydrolysis measurement.

Colorimetric assay was performed to analyze the EccA_1_ ATPase activity by using ATPase assay kit (*Innova Biosciences, UK*). The ATPase assay was performed in buffer containing (20mM HEPES pH 7.5, 150mM NaCl, 5% Glycerol, 2mM β-mercaptoethanol, 1mM ATP and 1mM of EccA_1_). The reaction was carried out at 25 °C for 5 min. The dye buffer containing (120mM malachite green, 0.06% polyvinylalcohol, 6mM ammonium heptamolybdate, 4.2% sodium citrate) was added in the reaction buffer. After incubating for 15 min, 10 μl of each reaction mixture was transferred in 96 well plates and absorbance at 630 nm was measured. Absorbance from reaction buffer containing no EccA_1_ and without ATP was substracted from each experimental data. Release of inorganic phosphate was analyzed based on absorbance from phosphate standard curve. Each assay was performed three times and average activity was calculated. The values of kinetic parameters *K*_m_ and *V*_max_ were calculated using PRISM 6.0 software (Graph Pad Software Inc.).

### Molecular modeling and dynamics simulation of EspC-EccA_1_ complex

PSIPRED [32] program was used to predict the secondary structures of EspC and EccA_1_. We obtained the EspC model by PHYRE2 server using WxG100 structure from *S. agalactia* (PDB-3GWK) as input. Full length EccA_1_ model (1-573 residues) was obtained by MODELER program [33]. Following templates were used as input (i) crystal structure of TPR domain of EccA_1_ (1-273 residues, PDB-4F3V) (ii) structural model of ATPase domain of EccA_1_ (331-481 residues) obtained by PHYRE2 server using Rubisco ATPase structure (PDB-3SYL) as input template [34]. Best model was selected based on MODELER Z-DOPE score. The EccA_1_ hexamer was built using COOT program [35] taking p97 hexamer (PDB-1E32) as input template [36]. Energy minimization was performed on EspC and EccA_1_ hexamer using GROMACS program[37]. PROCHECK program [38] was used to check the quality of EspC and EccA_1_ models and ANOLEA program [39] to calculate the conformational energy.

ROSETTADOCK server [40] with protein-protein docking module was used to build the EspC-EccA_1_ hexamer. The β□hairpin insertion motif (Ala81-Val96 residues) of TPR domain of EccA_1_ and YxxxD (Tyr87-Asp91 residues) motif of export arm of EspC were used as input constraints in protein-protein docking analysis. The docking server yielded the 10 best low energy clusters of EspC-EccA_1_ complexes. Best model was selected based on binding energy, intermolecular interactions and buried surface area in the EspC~EccA_1_ complex. The PISA server was used to analyze the interactions, binding free energy and solvent accessibility in EspC~EccA_1_ complex.

Energy minimization and dynamics simulation were performed on EspC~EccA_1_ complex using GROMACS program [37]. The EspC-EccA_1_ complex was immersed in cubic box extending 0.5 nm from protein surface and solvated with explicit SPC water molecules. The chloride and sodium ions were added to neutralize the whole system and simulated with periodic boundary conditions. The solvated complex system consists of 467,826 protein atoms surrounded by 1000,000 water molecules. Before running the dynamics simulation, whole system was energy minimized for 200 iterations of steepest descents and then equilibrated for 10ps, during which protein atoms were restrained. All restraints were removed from the protein and temperature was gradually increased in 10 distinct steps of 5 ps simulation each. V- rescale (modified Berendsen thermostat) coupling was employed to maintain a constant 300K temperature with a coupling constant of 0.2 ps. The coulomb cut off was 1.0. The time step employed was 2fs and coordinates are saved every 4ps for molecular dynamics trajectory analysis. The pressure was maintained by Parrinello-Rahman pressure coupling constant and coulomb cut off was applied during temperature coupling. The stereochemistry of simulated EspC~EccA_1_ complex was checked by PROCHECK program, secondary structure composition by DSSP program [41] and structure visualization by PyMOL [42] program.

### Binding analysis using wild type and mutant EspC and EccA_1_ proteins

Binding analysis between EspC and EccA_1_ was performed by surface plasmon resonance technique using BIAcore-3000 system (*Biacore Pharmacia Biosensor AB, Uppsala Sweden*). CM4 sensor chip was used for binding experiment and experiment was performed at 25°C in HBS-P buffer containing (20mM HEPES buffer pH 7.5, 150mM NaCl, 10mM Arginine, 10mM Glutamate, 0.02mM ATP, 0.02mM MgCl_2_). The CM4 sensor chip contains carboxy-methylated dextran covalently attached to gold surface and protein was covalently coupled to sensor surface via amine, thiol, aldehyde or carboxyl groups.

The EccA_1_ was immobilized on CM4 surface activated by EDC-NHS amine coupling chemistry. 170 μl of EccA_1_ (conc. ~ 60 μg/ml) was diluted in sodium acetate buffer pH 4.1 and injected over flow cells with 30 μl/min. The 5100 RU of EccA_1_was immobilized on CM4 sensor chip surface and surface was blocked by ethanolamine. Four different concentrations of EspC (0.5, 1, 2, and 4 μM) were injected over immobilized EccA_1_ using HBS-P buffer pH 7.5 with 30 μl/min. The sensogram was allowed to run for another 5 min. The biosensor surface was regenerated for another 10 min by using running buffer at 30μl/min. Association and dissociation kinetic constants were evaluated by BIAeveluation 3.0 software using simple 1:1 Langmuir model.

PCR based site directed mutagenesis was performed to obtain three deletion mutants of EspC, Surface plasmon resonance technique was used to obtain dissociation constant (*K_D_*) between EccA_1_ and three truncated mutants of EspC, (i) EspC-Δ5 (Met1-Asp99 residues) (ii) EspC-Δ10 (Met1-Trp93 residues) and (iii) EspC-Δ20 (Met1-Ala83 residues). To probe residues of EspC export arm involved in binding to β-hairpin insertion of TPR domain of EccA_1_, the EspC-W94A and EspC-Y87A mutants were generated. Binding analysis between five EspC mutants with EccA_1_were performed by same protocol used for EspC~EccA_1_ binding.Site directed mutants of EccA_1_ (D91A and Y89A in β-hairpin insertion of TPR domain) were generated and binding analysis was performed with EspC. The *K_D_* of two EccA_1_ mutants were obtained and compared with *K_D_* value obtained for EccA_1_~EspC binding.

### Circular dichorism and thermal stability analysis on wild type and mutant EspC and EccA_1_ proteins

CD measurements were recorded using Chirascan™ CD spectropolarimeter (*Applied Photophysics*), which uses the water bath to maintain the constant temperature. The wild type and mutant EspC and EccA_1_ proteins were diluted to 0.1 mg/ml in 10mM potassium phosphate buffer pH 8.0 and loaded in 0.1 cm quartz cuvette. The 10mM potassium phosphate buffer pH 8.0 was used as blank for all CD measurments. The final CD spectrum was average of three sequential scan. All CD data were converted to mean residue ellipticity (deg.cm^2^/dmol).

The Dichroweb server [43] was used to estimate the secondary structures from CD spectra. The SOPMA [44], GOR [45] and PSIPRED programs were used to estimate theoretical secondary structures of EspC and EccA_1_ proteins. For thermal stability analysis of EspC and EccA_1_, CD-spectra were recorded as a function of temperature. For EccA_1_, the CD spectra were recorded from 20°C - 90°C in 10°C increment and 25°C - 85°C in 10°C increment for EspC respectively.

## RESULTS

### EspC purification and characterization

The EspC protein (112 residues, Mw~11.9kD) contains a C-terminal export arm (with AA cradle, YSEADE and IDGLF motifs) involved in recognition to EccA_1_ ATPase (Figure 1A). The EspC was overexpressed in *E. coli. BL21(DE3)* cells and purified using Ni-NTA affinity and size exclusion chromatography (Figure 1B). Initially, the EspC was forming highly ordered oligomer and eluted in the void volume of Superdex 200(16/60) column. However, treatment with different additives improved the EspC solubility. Addition of 0.01%(w/v) Tween-20 and 0.01%(w/v) SDS in EspC buffer prevented the EspC oligomerization and protein eluted as hexamer on Superdex 200(16/60) column (Figure 1B). The mass spectrometric analysis on purified EspC (Figure S1) shows a single band and peak of EspC monomer (Mw~ 11.8 kD). The EspC was purified to homogeneity and found biologically active based on A549 cytotoxicity assay and binding analysis with EccA_1_ ATPase.

**Figure 1.**
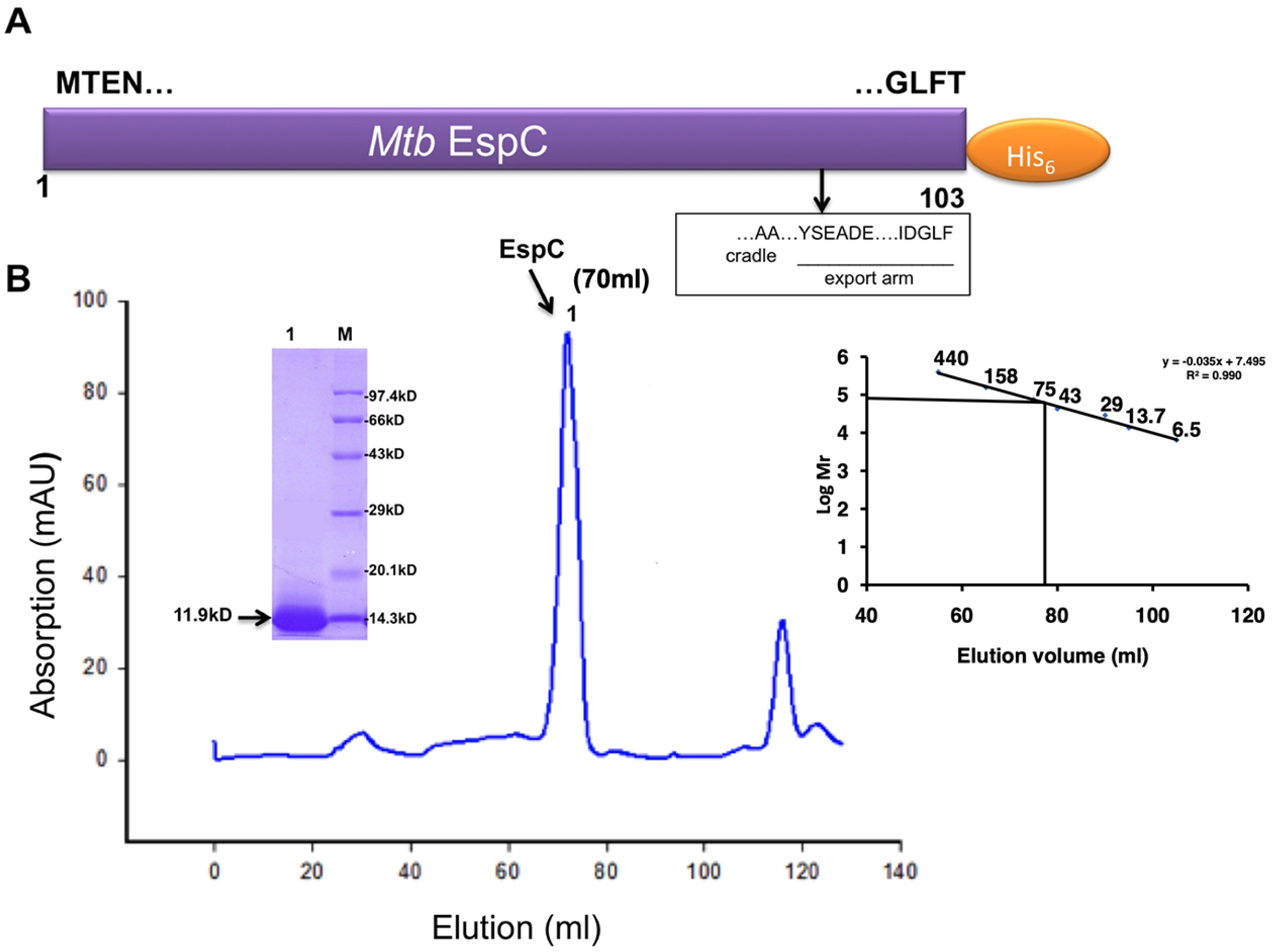
**A,** The schematic view of EspC and its expression construct. The EspC export arm consists of AA cradle, YSEADE motif and IDGLF motif. **B,** Size exclusion chromatography and SDS-PAGE analysis of purified EspC. EspC eluted as hexamer from Superdex200 (16/60) column and shows a single band on SDS-PAGE.

### EspC is cytotoxic to human lung carcinoma A549 cells

Human macrophage and epithelial cells constitute the first line of defense after mycobacterial infection and induce the initial host defense response. The human lung carcinoma A549 is a long epithelial cell that protects the host from pathogen infection. To check whether EspC is cytotoxic to A549 cell, we have performed the EspC cytotoxicity experiment using A549 cells and examined the results using light microscopy and MTT assay.

As seen in MTT assay (Figure 2), viability of A549 cells decreased upon incubation with EspC protein. Four different concentrations of EspC (25, 50, 100 and 200 *μ*g/ml) were incubated with A549 cells for different time points (2, 4, 6, 12 and 24 h). Viability of A549 cells decreased progressively with increasing concentration of EspC. There was significant decrease in A549 cells viability with EspC conc. of 100 and 200 *μ*g/ml, when compared with Bovine Serum Albumin in EspC buffer as a control (Figure 2). With 100 *μ*g/ml of EspC at 2, 4, 6, 12 and 24 h, % of cell viability (53.9, 41.8, 37.5, 17.6, 4.73) was observed. With 200 μg/ml of EspC, % of cell viability (4.4, 1.46, 1.36, 1.23, 0.75) was observed. Phospholipase-C was used as a positive control, which resulted significant decrease in A549 cells viability at all time points used in the study. Bovine Serum Albumin (Conc. ~ 100 *μ*g/ml dissolved in EspC buffer) did not show any significant decrease in cell viability (Figure 2). These results indicate that EspC is cytotoxic to the human lung carcinoma A549 cells.

**Figure 2.**
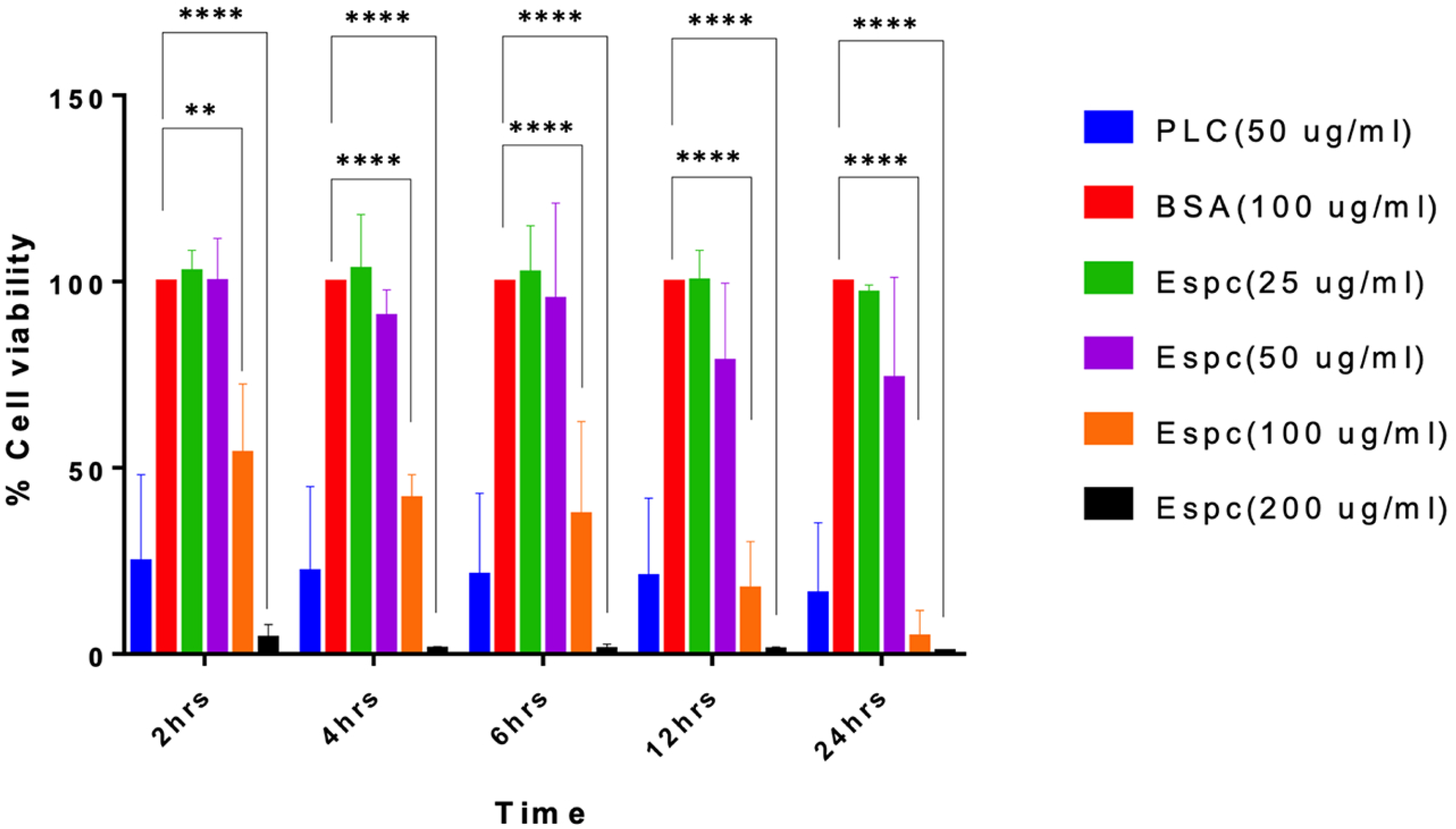
The purified EspC is cytotoxic to A549 cells. MTT assay was performed to evaluate the cytotoxic potential of EspC in A549 cells. A549 cells were treated with the EspC (Conc.~ 20, 50, 100 and 200 μg/ml) for 2, 4, 6, 12 and 24 h. The BSA (Bovine serum albumin) dissolved in EspC-buffer containing detergents was used as negative control. The phospholipase-C (PLC) in EspC buffer was used as positive control for cytotoxic lysis experiment. After incubation, the A549 cells were harvested and treated with MTT (conc. ~ 5 mg/ml). The Live cells and dead cells using MTT assay were counted at different conditions. The graph shows total number of live cells (Y-axis) plotted for different time durations (X-axis) under different conditions. Ns-non significant, ***P<0.001, ****P<0.0001.

### EspC enters into A549 cells

To examine, whether EspC enters into A549 cells and indeed cytosolic, we stained the EspC (100 *μ*g/ml) treated A549 cells with anti-His antibody and analyzed the cells using confocal microscopy (Figure 3). We examined the localization of EspC in A549 cells at 2, 4 and 6 h using confocal microscopy, as these time points resulted in equally significant decrease in cell viability as the later time-points. The confocal images show that EspC was localized in the cytosol of A549 cells after 2, 4 and 6 h of treatment. The A549 cells without EspC treatment did not stain with anti-His antibody. These results show that EspC enters into A549 cells and results in decreased cell viability.

**Figure 3.**
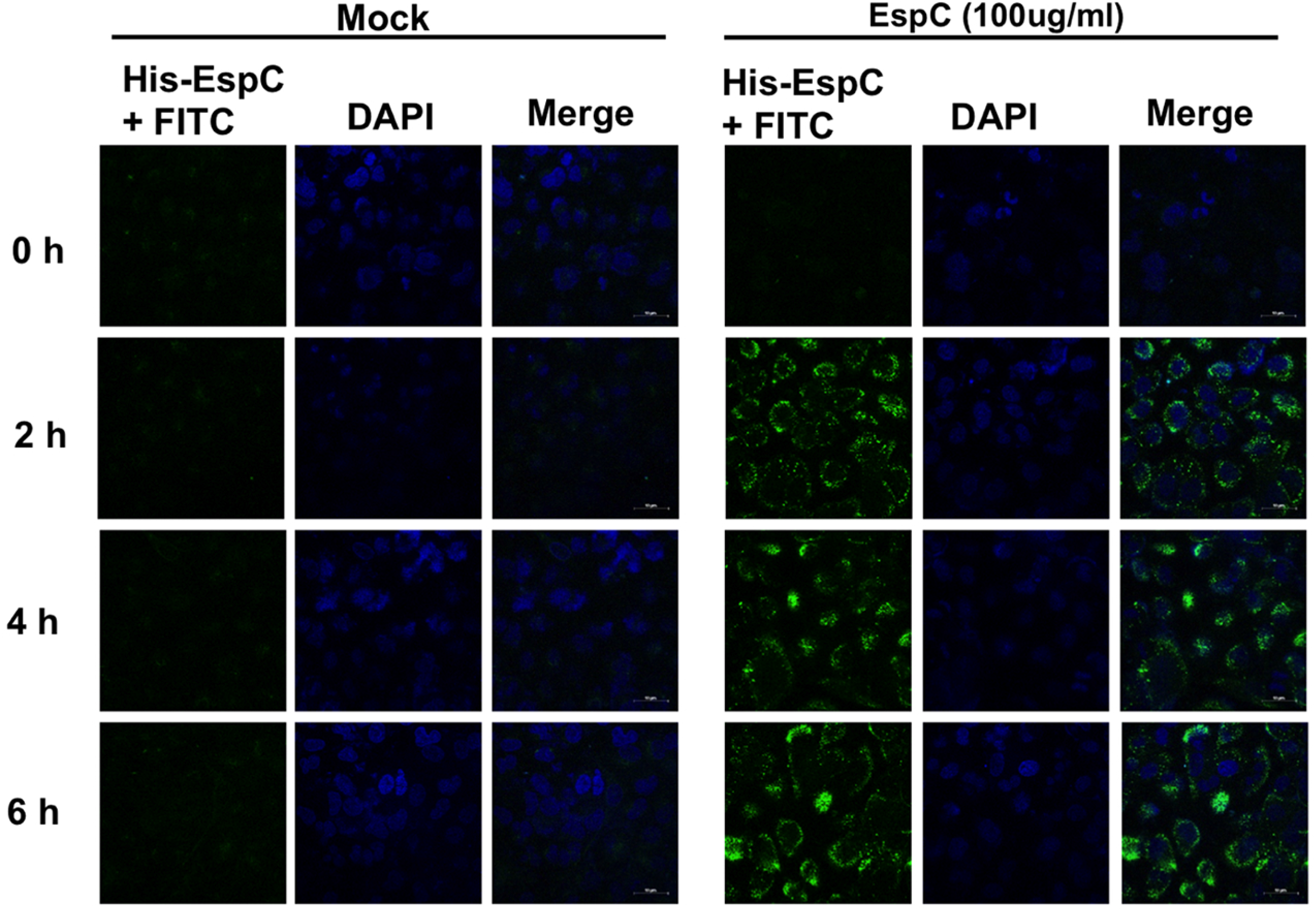
The EspC protein is localized intracellular in the A549 cells. Representative images of confocal microscopy showing the EspC accumulation at 0, 2, 4 and 6 h post treatment with EspC protein. A549 cells were seeded on coverslip and after 24 h cells were treated with EspC-buffer (mock) and with Esp Cprotein(conc. ~100 μg/ml)for 2, 4, 6 h. The presence of EspC protein (green) was tested with anti-His antibody, as EspCwas tagged with 6xHis. DAPI and Merge (His-EspC + DAPI) images are shown. White scale bar is 10μm.

To test the amount of EspC present in A549 cells, the EspC was incubated for 2, 4, and 6 h with A549 cells and presence of EspC in whole cell lysate was detected using anti-His antibody (Figure 4C). Specific bands corresponding to EspC were detected only in EspC treated A549 cell lysate and not in control experiment. The EspC band was detected at 2, 4 and 6 h of treatment of A549 cells with EspC.

**Figure 4.**
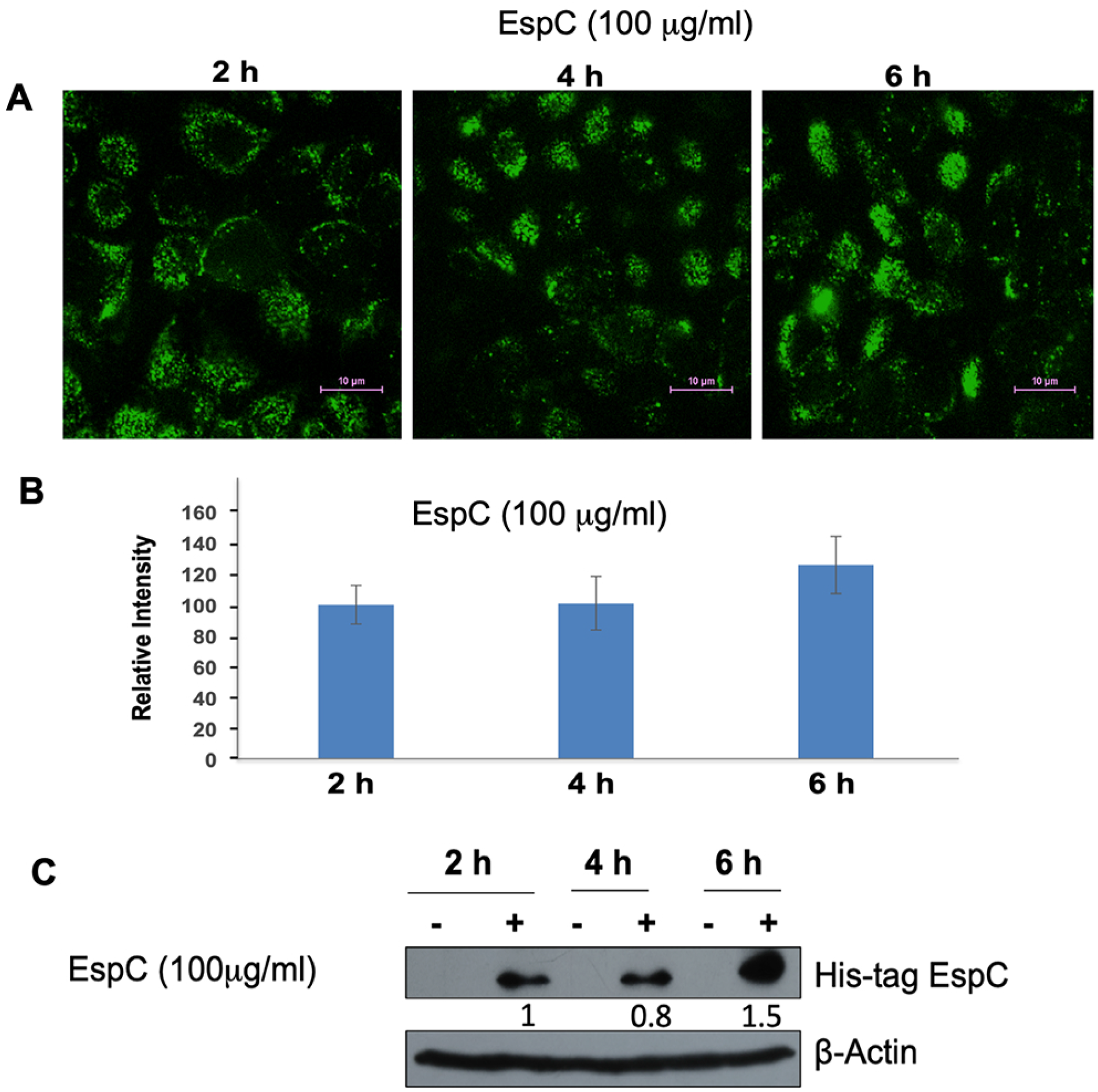
**A,** The EspC accumulates inside the A549 cells gradually with time. Representative images show the EspC protein (100 μg/ml, green) treated with A549 cells for different time points, 2, 4 and 6 h. **B,** Quantitative estimation of fluorescence intensity at each time was performed using Image J software [54]. The relative intensity (Y-axis) was plotted for different time points (X-axis). The graph represents the data from two independent biological experiments performed in duplicates. The error bars represent standard deviation in two experiments. White scale bar is 10μm. **C,** The EspC enters and localize inside the A549 cells. The EspC protein (conc. ~ 100 μg/ml) was incubated for different time points (2, 4, and 6 h) with A549 cells. The treated cells were tested for EspC presence inside the A549 cells by Western blot analysis. A blot probed with anti His-antibody (for detection of EspC protein) showed the accumulation of EspC protein after 2, 4 and 6 h (+). The negative control (-) (no EspC incubation) did not show presence of EspC. A parallel blot probed with anti β-actin antibody is shown as a loading control. Numbers between His-tagged EspC and β-actin denote the relative intensity of EspC band.

### EccA_1_ shows the specific ATPase activity

The schematic view of EccA_1_ ATPase (67kD, 573 residues) is shown in Figure S2A. The The EccA_1_ protein was eluted as hexamer from Superdex 200(16/60) column and showed a single band on SDS-PAGE (Figure S2B). Initially, EccA_1_ was eluting in the void volume of Superdex 200(16/60) column, however addition of 10mM Arginine, 0.2mM MgCl_2_ and 0.2mM ATP in protein buffer prevented the EccA_1_ oligomerization and protein eluted as hexamer. The AAA+ATPase proteins may form multiprotein complex inside the cell and higher ordered oligomers of EccA_1_ hexamer have been observed in earlier study also [46].

To examine the ATPase activity, EccA_1_ was dialyzed in buffer containing no ATP. We performed sensitive radioactive assay on EccA_1_ using [γ^32P^]ATP as a substrate and release of free phosphate was monitored (Figure S2A). The free phosphate was released linearly overtime and *K_m_* of 52.4±2.1 μm and *V*_max_ of 1.51±0.7 μmol/min were obtained from Michaelis-Menten plot and nonlinear regression analysis using Prism 6.0 software (GraphPad Software) (Figure S2B). These data shows that EccA_1_ is a weak ATPase, as also observed in earlier study [46].

### Molecular modeling of EspC-EccA_1_ complex

The sequence alignment of the export arm of EspC with other ESX-1 virulence proteins is shown in Figure 5A. The EspC may form homodimer, as observed for monocistronically expressed CFP10 and sagEsxA family proteins, which lack WxG motif (24). The model of EspC homodimer (Figure 5B) was obtained by PHYRE2 server (47), which used the sagEsxA structure (PDB-3GWK) as input template. As EspC eluted as hexamer from gel filtration column, we build the model of EspC hexamer by ROSETTADOCK server (40) using Cn symmetry module (Figure S4). The lowest energy cluster was selected as the final model, which shows the export arms of the EspC on both sides of helical bundle and easily accessible for optimal interactions with EccA_1_.

**Figure 5.**
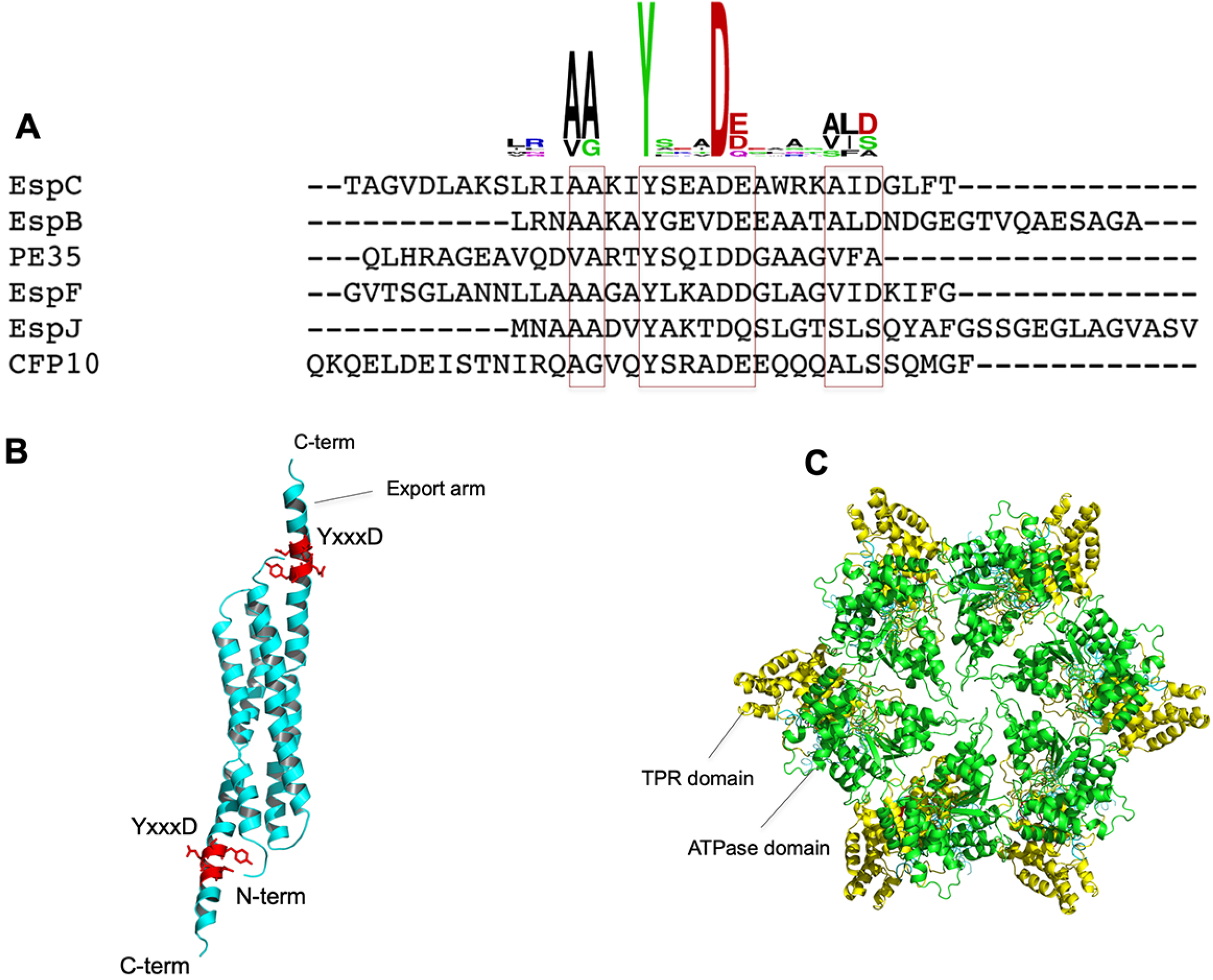
**A,** Sequence alignment of export arm of EspC with other ESX-1 virulence proteins. The conserved AA-cradle, YxxxD motif and ALD motif of the EspC protein are shown in red bracket. The Web-LOGO diagram is shown above the sequence alignment. **B,** The EspC dimer is modeled using SagEsxA structure as input template. The red color indicates the YxxxD motif of the export arm of EspC, involved in binding to EccA_1_. **C,** The EccA_1_ hexamer model is obtained using the Modeler program. The yellow color indicates the N-terminal TPR domain, and green color shows the C-terminal ATPase domain of the EccA_1_.

The EccA_1_ model was obtained using following constraints. The model of C-terminal ATPase domain (331-481 residues) of EccA_1_ was obtained using Rubisco ATPase structure (PDB: 3ZUH) as input. The structure of the N-terminal TPR domain (1-273 residues) was obtained from published structure of *M. tuberculosis* TPR domain (PDB: 4F3V) (31). The structure of full length EccA_1_ (1-573 residues) was obtained by MODELER program [30, 36] using following structures, the N-terminal TPR domain, the C-terminal hexameric ATPase model and two loops connecting 273-331 residues and 481-573 residues respectively. The EccA1 hexamer was built using p97 hexamer structure (PDB-1E32) as input model (Figure 5C).

To build EspC-EccA_1_ complex, ROSETTADOCK protein-protein docking server was used with following constraints (i) the β-hairpin motif (A81-V96 residues) of the TPR domain of EccA_1_ and YxxxD motif (Y87-D91 residues) of EspC homodimer as interacting partner in protein-protein docking calculation. Simulations with stoichiometric constraints resulted convergence to several low energy clusters of EspC-EccA_1_ complex. All low energy conformations have similar binding mode in EspC-EccA_1_ complex and that provides greater confidence in proposed orientation of both molecules in EspC-EccA_1_ complex. The lowest energy cluster was selected and dynamics simulation was performed on EspC-EccA_1_ complex using GROMACS program [37] (Figure 6A-D).

**Figure 6.**
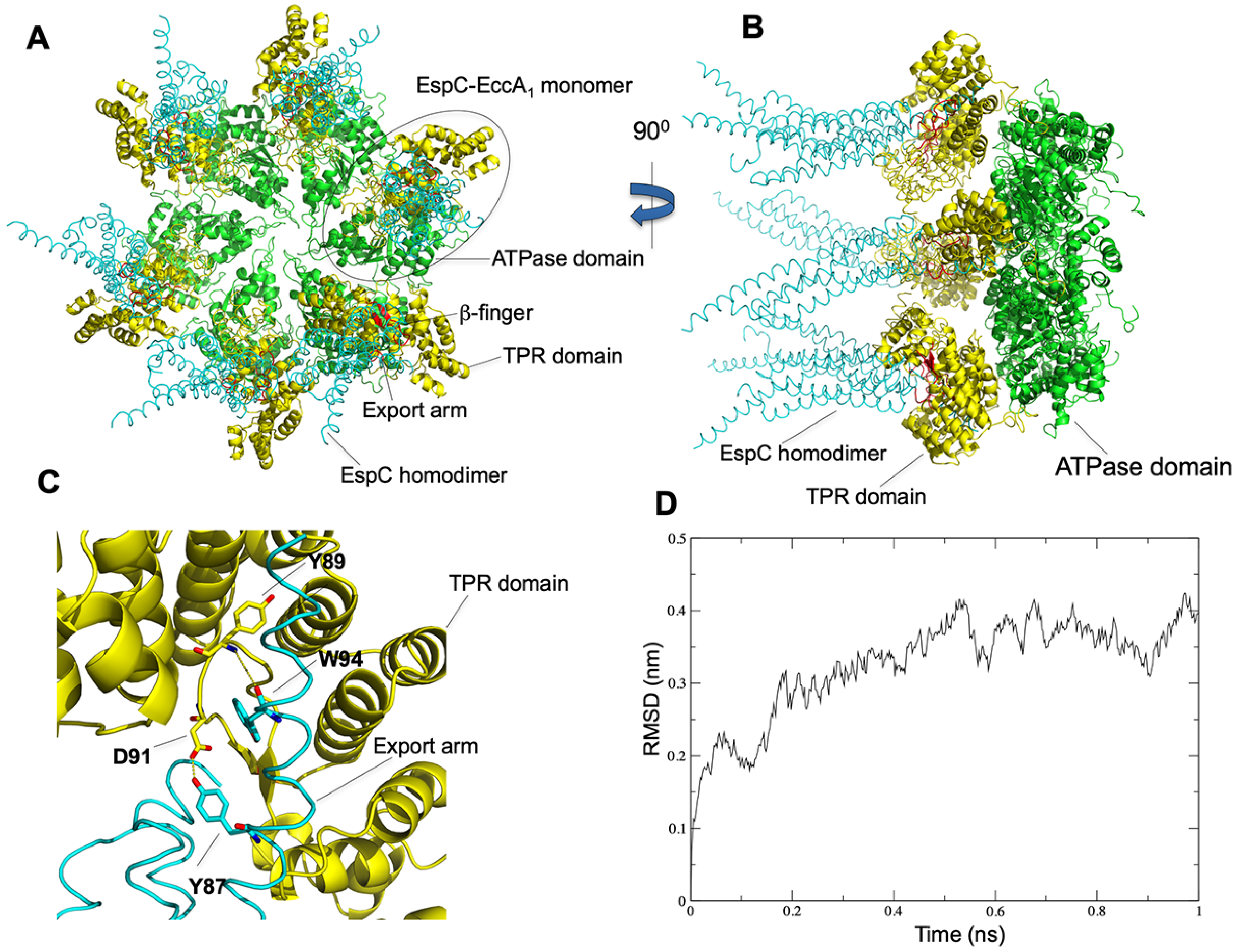
**A,** Homology modeling, protein-protein docking and dynamics simulation were used to obtain the EspC-EccA_1_ complex model. The EspC homodimer (cyan), TPR domain (yellow) and ATPase domain are shown in green color. β-hairpin insertions of TPR domain of EccA1 and export arm of EspC are shown in red color. **B,** The EspC-EccA_1_ complex view after 90^0^ of rotation**. C,** closer view of interactions between EspC exports arm and β-hairpin insertion of TPR domain of EccA_1_. **D,** Root mean square deviation of Cα atoms during dynamics simulation with respect to the starting structure.

PISA server [48] was used to analyze the interface between EspC and EccA_1_ proteins. It yielded following hydrogen bonds and salt bridges at EspC~EccA_1_ interface, (i) D91(Oδ1) of EccA_1_ with Y87(N) of EspC (ii) Y89(OH) of EccA_1_ with W94(Nζ) of EspC (iii) A112(O’) of EccA_1_ with A93(N) of EspC (iv) Y16(OH) of EccA_1_ with Ala93(O’) of EspC (v) R206(Nε) of EccA_1_ with Y87(OH) of EspC (vi) E238(Oε1) of EccA_1_ with Y87(OH) of EspC and (viii) T234(O’) of EccA_1_with R11(NH_1_) of EspC. These data indicate that EspC export arm interacts extensively with β–hairpin insertion of the TPR domain and N-terminal regions of the EccA_1_ ATPase.

### EspC has a bipartite signal sequence that interacts with TPR domain of EccA_1_

The C-terminal export arm of EspC harbors the AA-cradle (A94-A95), YSEAD motif (Y87-D91) and IDGLF (I98-F102) motifs. The EspC export arm interacts with EccA_1_ ATPase and required for EspC secretion by ESX-1 system [30]. To identify the residues and length of EspC export arm involved in EccA_1_ recognition, we prepared three truncated (EspC-Δ5, EspC-Δ10 and EspC-Δ20) and two point mutants (W94A and Y89A) of EspC export arm, involved in binding to TPR domain of EccA_1_.

The surface plasmon resonance technique was used for binding analysis between EspC and EccA_1_, which showed the *K_D_* value of ~ 0.14μM±0.05μM (Figure 7A). From EspC-EccA_1_ complex model, we identified that EspC-Y87 residue forms hydrogen bond with D91 residue of β-hairpin insertion of TPR domain. The EspC-W94 residue forms hydrogen bond with Y89 residue of β-hairpin insertion of the TPR domain of EccA_1_. We performed the binding analysis of EspC-W94A mutant with EccA_1_ and obtained *K_D_* value of 9.1±0.8μM (Figure 7B) and 10.3±0.7μM for EspC-Y87A mutant (Figure 7C). ~ 65 fold reduction in binding affinity for EspC-W94A mutant and ~ 74 fold reduction in binding affinity for EspC-Y87A mutant were observed, when compared with affinity between EspC~EccA_1_. These results indicate that W94 and Y87 residues of EspC are critical in recognition to TPR domain of EccA_1_.

**Figure 7.**
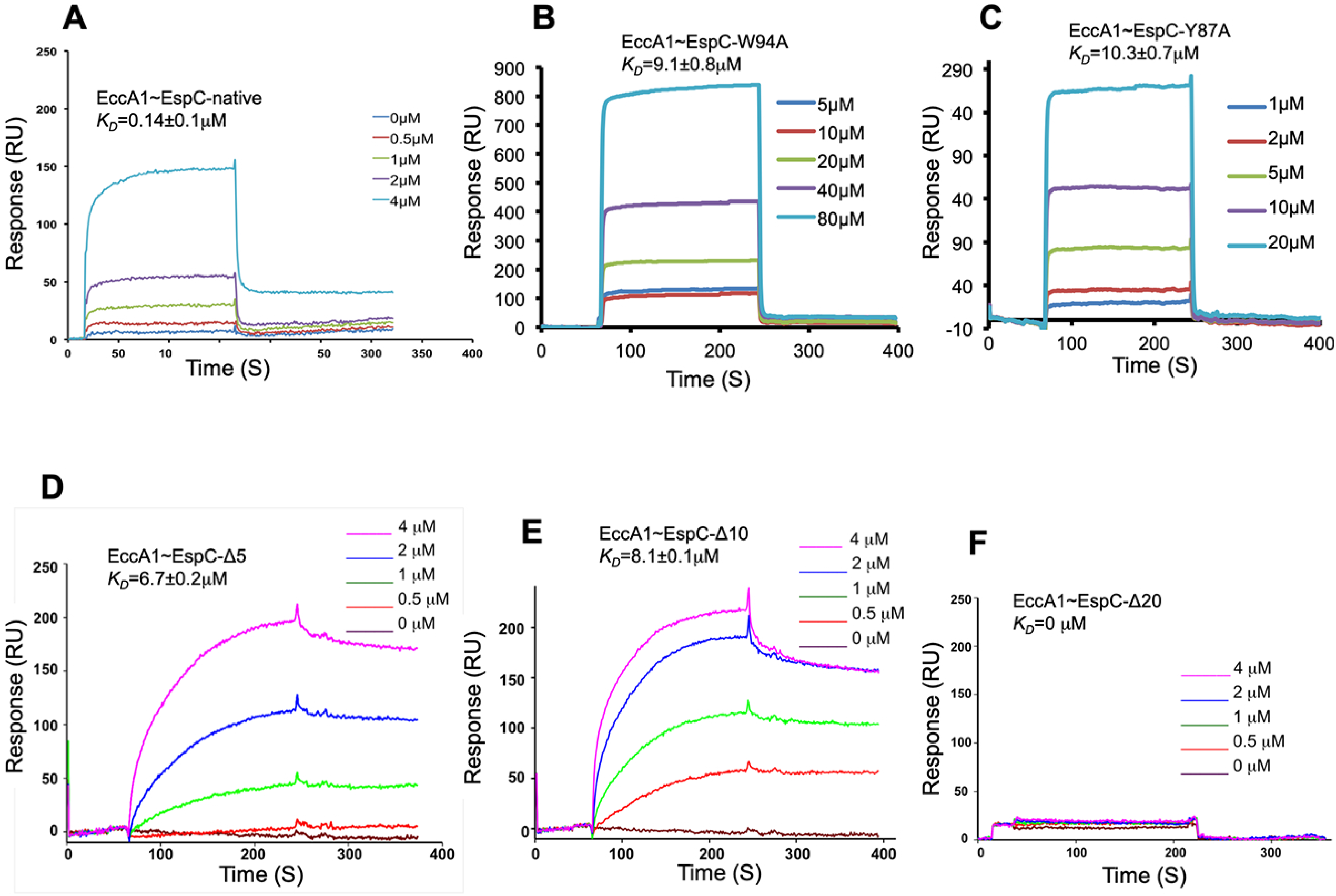
The binding analysis of EccA_1_ was performed with five EspC mutants using surface plasmon resonance technique. **A,** Binding analysis of EccA_1_ with EspC. **B,** Binding analysis of EccA_1_with EspC-W94A mutant. **C,** Binding analysis of EccA_1_ with EspC-Y87A mutant. **D,** Binding analysis of EccA_1_ with EspC-Δ5 mutant. **E,** Binding analysis of EccA_1_ with EspC-Δ10 mutant. **F,** Binding analysis of EccA_1_ with EspC-Δ20 mutant. The concentrations of EspC mutants used in EccA_1_ binding and obtained *K_D_* values are shown in the each figure.

To probe the length of EspC export arm involved in EccA_1_ recognition, we generated three truncated mutants (i) EspC-Δ5 (ii) EspC-Δ10 and (iii) EspC-Δ20. The EspC-Δ5 and EspC-Δ10 mutants lack conserved IDGLF motif of the export arm and EspC-Δ20 mutant lacks full export arm (AA-cradle, YSEAD motif and IDGLF motif). Binding analysis of wild type EccA_1_ with three truncated mutants of the EspC yielded following *K_D_* values, (i) EspC-Δ5 (*K*_D_~6.7±0.2 μM) (Figure 7D) (ii) EspC-Δ10 (*K*_D_~8.1±0.1 μM) (Figure 7E) and (iii) EspC-Δ20 (no binding) (Figure 7F). These results indicate that EspC-Δ5 mutant (lacking IDGLF motif) showed ~ 40 fold and EspC-Δ10 mutant (lacking WRKA-IDGLFT motif) shows ~ 60 fold decrease in binding affinity, when compared with wild type EspC affinity towards EccA_1_. The EspC-Δ20 mutant (lacking full export arm consists of AA-cradle, YSEAD and IDGLF motifs) showed ~ no binding to EccA_1_. Altogether, these results indicate that IDGLF motif of EspC export arm is critical for EccA_1_ binding and its deletion in EspC-Δ5 and EspC-Δ10 mutants leads to ~ 40-60 fold decrease in binding to EccA_1_. Conserved AA-cradle, YSEAD and IDGLF motifs of EspC export arm are absolutely essential for EccA_1_ recognition and deletion of these motifs in EspC-Δ20 mutant leads to ~ no binding to EccA_1_.

### β□hairpin insertion motif of TPR domain of EccA_1_ interacts specifically with export arm of EspC

Based on EspC-EccA_1_ model, two point mutations (Y89A and D91A) in β-hairpin insertion motif of TPR domain were generated using site directed mutagenesis. D91 residue of β-hairpin insertion forms hydrogen bond with Y87 residue of EspC export arm (^87^YxxxD^91^). The Y89 residue of β-hairpin insertion forms hydrogen bond with W94 residue of EspC export arm. Binding analysis of EspC with wild type & two EccA_1_ mutants was performed using surface plasmon resonance technique. It shows that wild type EccA_1_ binds to EspC with *K_D_* ~0.14 μM (Figure 8A). The EccA_1_-D91A mutant binds to EspC with *K_D_* ~ 3.2 μM (Figure 8B) and EccA_1_ -Y89A mutant binds to EspC with *K_D_* ~2.6 μM (Figure 8C). These data indicate that EccA_1_ -Y89A mutation leads to ~19 fold decrease and EccA_1_-D91A mutation leads to ~23 fold decrease in binding affinity to EspC, when compared with wild type EccA_1_. These results show that Y89 and D91 residues of β-hairpin insertion motif of TPR domain specifically recognize the EspC export arm and their mutations lead to significant decrease in EspC binding.

**Figure 8.**
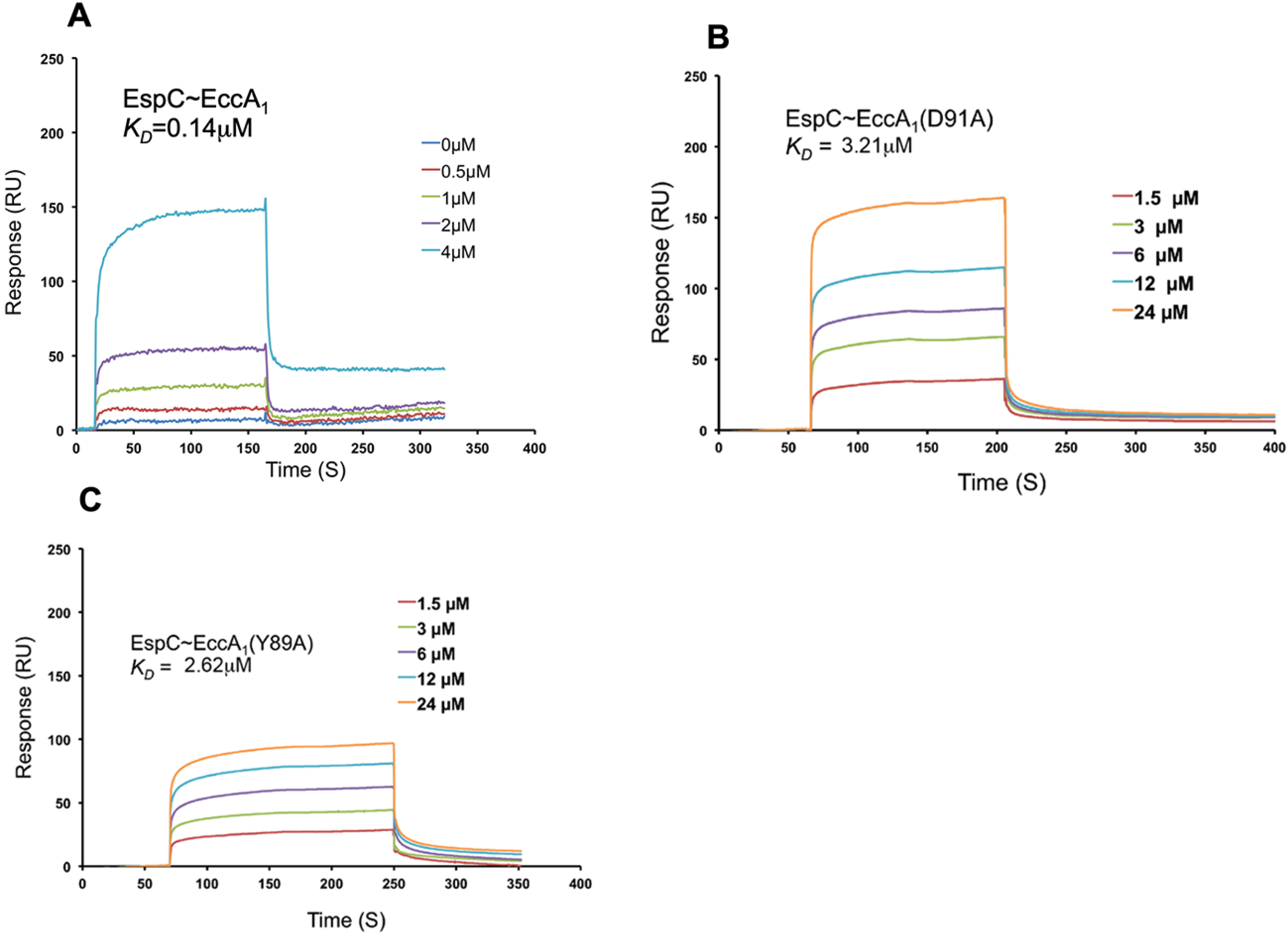
**A,** Binding analysis of EspC with EccA_1_ mutants using surface plasmon resonance technique. **A,** Binding analysis of EspC with EccA_1_. **B,** Binding analysis of EspC with EccA_1_-D91A mutant. **C,** Binding analysis of EspC and EccA_1_-Y89A mutant. The concentrations of the EccA_1_ mutants used in EspC binding and obtained *K_D_* values are shown in each figure.

### The EspC and EccA_1_ show quite similar secondary structures in wild and mutant proteins respectively

Far-UV CD spectra (260-200 nm) were obtained for native EspC, EspC-W94A, EspC-Y87A, EspC-Δ5, EspC-Δ10, and EspC-Δ20 mutants (Figure 9A-F). Dichroweb server was used to estimate the secondary structures and thermal stability of all proteins (Table 1) [43]. As seen from Table 1, the wild type EspC contains ~ 80% α-helix and ~ 20% random coil structures, which was quite similar to secondary structures in five EspC mutants. These results show that wild type and mutant EspC proteins retain quite similar structures. For thermal stability analysis, the CD spectra of EspC were recorded from 25°C - 85°C in 10°C increment (Figure 9A). These data shows the thermal unfolding transition temperature (T_m_) of 32.2±1.0 °C, which indicate that EspC is a quite thermostable protein.

**Figure 9.**
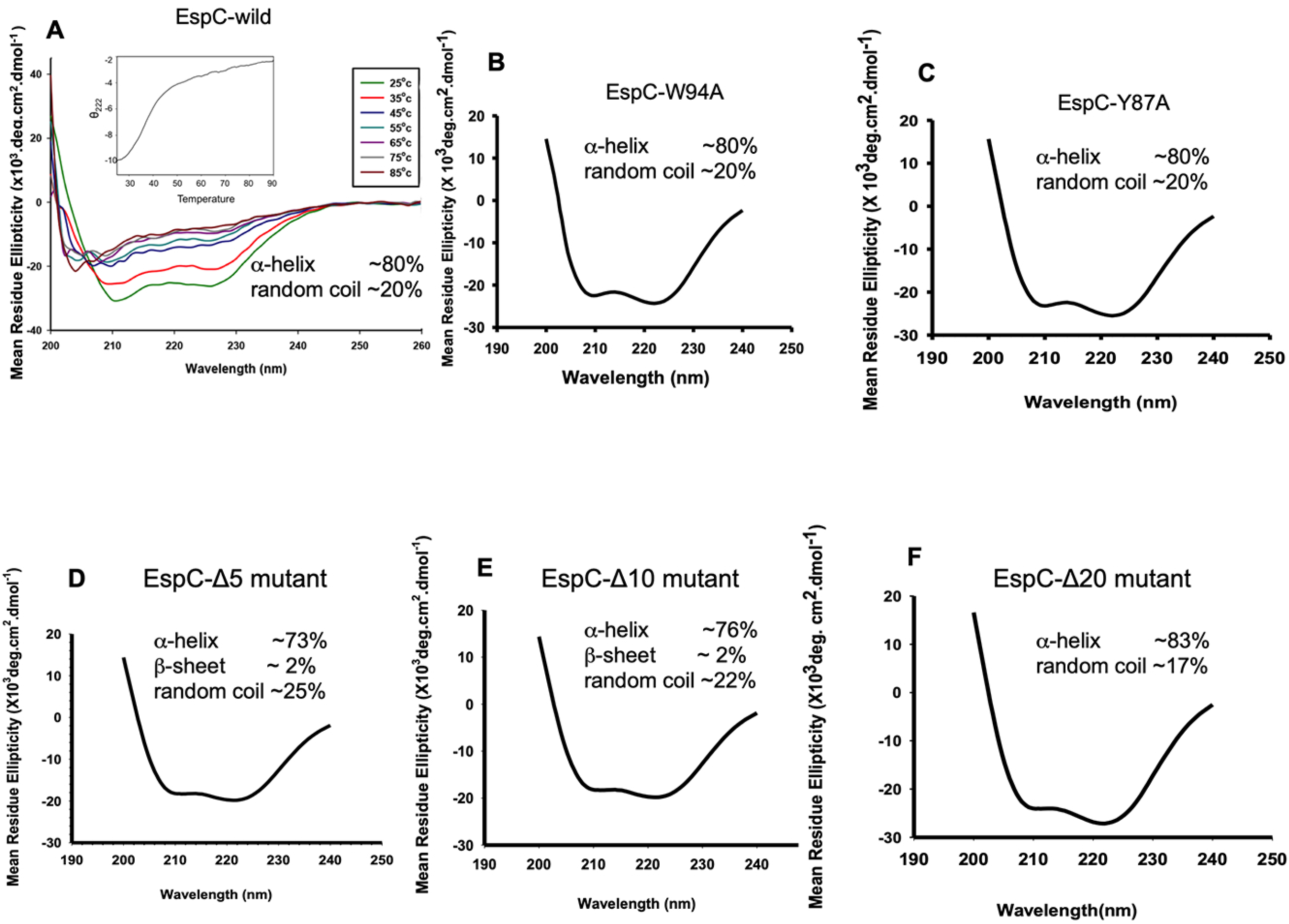
**A,** Circular dichorism and thermal stability profile of EspC. **B-C,** Circular dichorism spectra of EspC-W94A and EspC-Y87A mutants. **D-F,** Circular dichorism spectra of EspC-Δ5, EspC-Δ10 and EspC-Δ20 mutants. Secondary structures of wild type and mutants EspC were determined using Dichroweb server and yielded quite similar values, as shown in each figure. Thermal stability analysis on EspC shows the T_m_~ 32.1 °C

**Table 1:**
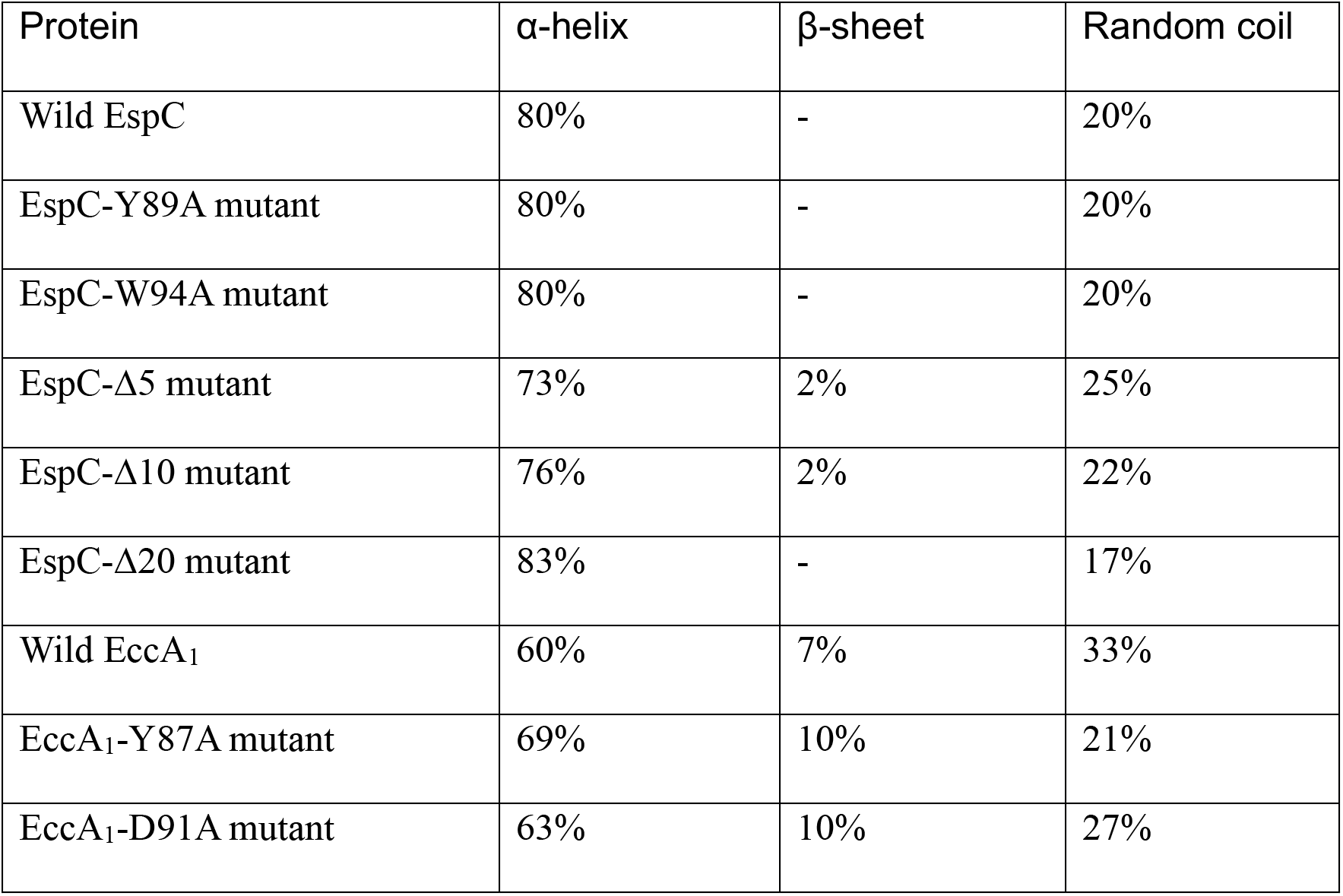
Circular dichorism analysis of wild and mutant EspC and EccA_1_ proteins

| Protein | α-helix | β-sheet | Random coil |
| --- | --- | --- | --- |
| Wild EspC | 80% | - | 20% |
| EspC-Y89A mutant | 80% | - | 20% |
| EspC-W94A mutant | 80% | - | 20% |
| EspC-Δ5 mutant | 73% | 2% | 25% |
| EspC-Δ10 mutant | 76% | 2% | 22% |
| EspC-Δ20 mutant | 83% | - | 17% |
| Wild EccA<sub>1</sub> | 60% | 7% | 33% |
| EccA<sub>1</sub>-Y87A mutant | 69% | 10% | 21% |
| EccA<sub>1</sub>-D91A mutant | 63% | 10% | 27% |

Far-UV CD spectra (260-200 nm) were obtained for wild type EccA_1_, EccA_1_-D91A and EccA_1_-Y87A mutants (Fig 10A-D). Dichroweb server analysis on wild type and two EccA_1_ mutants shows quite similar secondary structures (~ 60% α-helix, ~ 7% β–sheet and ~ 33% random coil structures) (Table 1). The theoretical structure prediction analysis on EccA_1_ also shows the ~ 60% α-helix, ~ 4.3% β–sheet, ~ 35.2% random coil structures, which was quite similar to CD data. For thermal stability analysis, the CD spectrum for EccA_1_ was collected from 20°C – 90°C in 10°C increment (Figure 10B). It shows the thermal unfolding transition temperature (T_m_) of 49.3±0.1 °C, which indicate that EccA_1_ is a quite thermostable protein.

**Figure 10.**
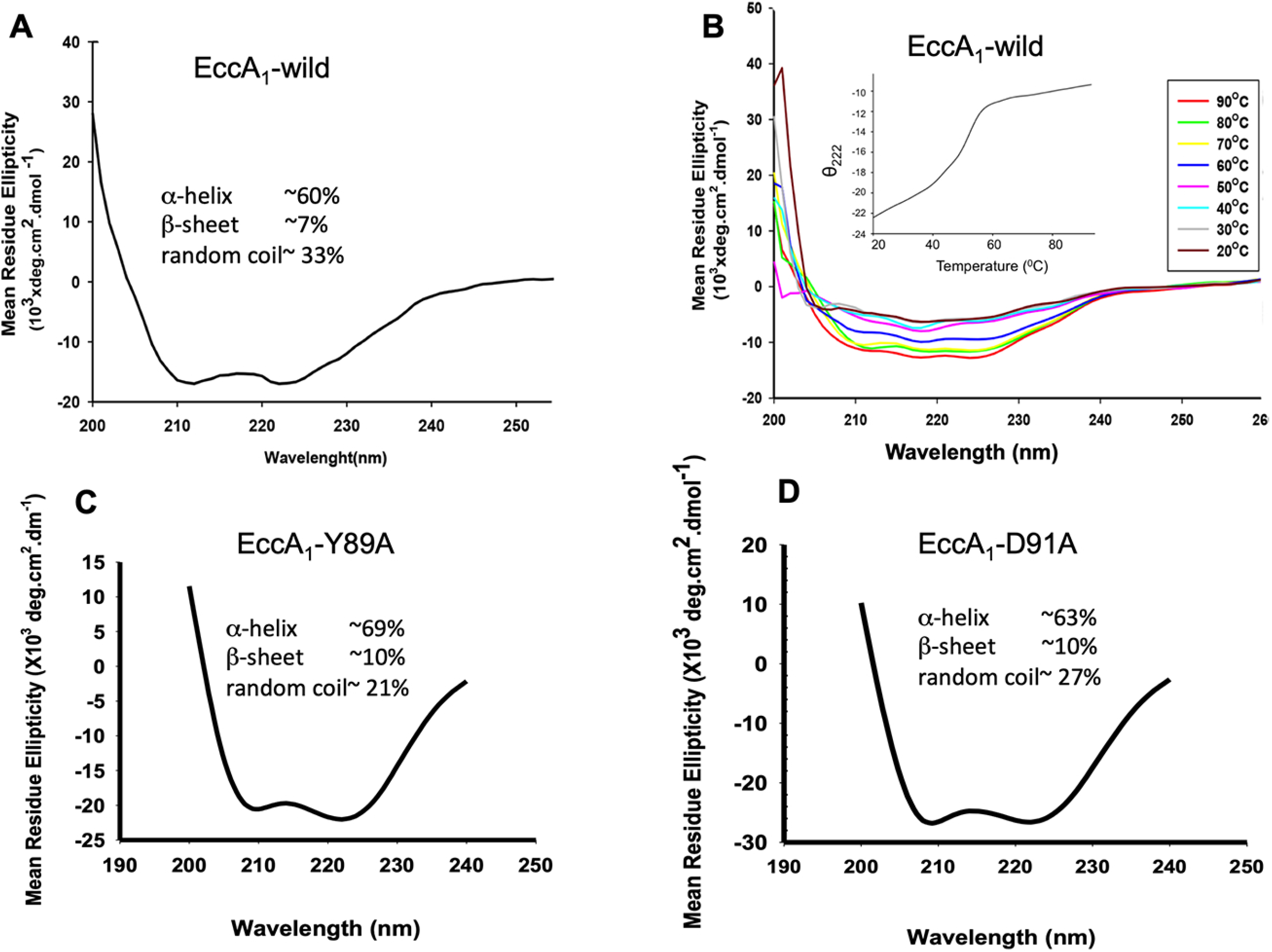
A-B, Circular dichorism and thermal stability spectra of the EccA_1_. C-D, Circular dichorism spectra of EccA_1_-Y89A and EccA_1_-D91A mutants. Spectra were recorded at room temperature and normalized to mean residue ellipticity.

## DISCUSSION

In the current study, we have purified and analyzed the biological activities of EspC and EccA_1_ proteins. To understand the molecular mechanism involved in EspC~EccA_1_ recognition, molecular modeling and docking techniques were used to obtain the EspC~EccA_1_ model. Based on modeled structure, critical residues involved in EspC ~ EccA_1_ recognition were identified and EspC and EccA_1_ mutants were generated. To validate these interactions, binding analysis was performed using wild type and mutant EspC to EccA_1_ proteins respectively to dissect the molecular mechanism involved in EspC~EccA_1_ recognition. CD spectroscopy was used to identify the secondary structure and thermal stability of EspC and EccA_1_ proteins as a result of all mutations.

We have expressed and purified the EspC protein in detergent solubilized form. The wild type and mutant EspC proteins eluted as hexamer from the size exclusion column. It is observed that EspC co-precipitates with EspA in cytosolic and membrane fractions of the mycobacterial cell [18]. In absence of EspA, the EspC may form homodimer as observed for monocistronically expressed CFP10 and sagEsxA family proteins, which lack WxG motif [24]. It appears that EspC hexamer may assemble as three copies of dimer in solution (Figure S4). The ESX-1 virulence proteins have been found to form multiprotein complex before their secretion by ESX-1 system.

The *Mycobacteria* uses the ESX-1 mediated membrane-puncturing mechanism for phagosomal permeabilization within infected host macrophage. By secreting EspC and other ESX-1 virulence proteins, mycobacteria may be able to escape phagosome and enter into the host cytosol. To examine whether EspC enters into the human lung carcinoma A549 cells and induce cytotoxicity, we incubated the EspC with A549 cells and examined the viability of the cells using MTT assay. The EspC cytotoxicity could be detected in A549 cells as early as 2 h of incubation and gradually increased. Contrary to our result, S. T. Cole group [18] has reported that they did not find any detectable cytotoxicity in RAW264.7 cells upon incubation with EspC for 2 h. The EspC enters into the A549 cells and accumulates in the cytosol, as observed in confocal microscopy and Western blotting analysis. Similar results were observed with *Listeria monocytogenes* bacterial proteins, which disrupt the vacuole membrane and help bacteria to enter in cytosol of host cell [40]. Significant cytotoxicity in A549 cells due to recombinant EspC was evident only at higher concentrations of EspC. This could be due to limited permeability of the recombinant EspC in A549 cells and hence lesser effective concentration inside the target cells under *in vitro* conditions. During natural infection of the target macrophages with *Mycobacterium*, the protein might be directly injected inside the cell and hence higher effective concentration.

The EccA_1_ belongs to CbxX/CfxQ family of ATPases, a classical AAA+ATPase protein family. The EccA_1_ forms hexamer through C-terminal ATPase domain and contains a unique N-terminal TPR domain. We have performed the ATPase assay using full length EccA_1_, which shows weaker ATPase activity compared to other AAA^+^ ATPases. In earlier study [46], fulllength EccA_1_ had shown weaker ATPase activity, when compared with ATPase activity of C-terminal domain alone. Other AAA^+^ ATPase enzymes also have shown weaker ATPase activity like *E. coli*. ClpB [47]. It is observed that loading of EspC on N-terminal TPR domain of EccA_1_ enhances the ATPase activity [46]. The ClpB ATPase activity is also enhanced in presence of casein [47], as observed in case of EccA_1_ protein.

The TPR domain uses its concave groove for protein-protein interaction and usually found in multi-protein complexes [48–51]. The EccA_1_ uses the concave groove of TPR domain for EspC binding, however a β-hairpin insertion in observed in the concave groove, which may form extended β-strand complementation with export arm of the EspC. The EspC may also interact on the side of concave groove of TPR domain as observed in P67-TPR/Rac complex structure [52]. To understand this mechanism, we have modeled the EspC~EccA_1_ complex and found that β-hairpin insertion of TPR domain interacts extensively with EspC export arm. Based on these interactions, we designed EspC and EccA_1_ mutants and performed binding analysis using wild type and mutant proteins. Mutations in EspC export arm/ or in β□hairpin insertion of TPR domain leads to significant reduction in binding affinity between EspC and EccA_1_ in comparison to wild type EspC~EccA_1_ affinity. To examine, whether wild type and mutant EspC and EccA_1_ proteins maintain the similar structures, we performed the CD spectroscopy and thermal stability analysis on wild type and mutant proteins. No significant changes in secondary structures were observed in mutant proteins, when compared with wild type EspC and EccA_1_ proteins.

These results have allowed us to rationalize the binding mechanism and specificity determinants at EspC~EccA_1_ interface. Biochemical results obtained from binding analysis of wild type and mutant proteins are in good agreement with interactions observed at EspC~EccA_1_ interface. Truncation/mutation in EspC export arm or point mutations in β-hairpin insertion of TPR domain of EccA_1_ caused significant decrease/or abolished the EspC~EccA_1_ binding. Interactions observed at EspC~EccA_1_ interface are quite similar to the interactions observed in Rac/p67-TPR domain complex. In Rac/p67-TPR domain complex, Rac binds p67-TPR concave groove in extended chain conformation and binding occurs exclusively with β–hairpin insertion. TPR domains are quite versatile in target recognition and follow the general rule for scaffolding the multiprotein complex assembly as observed in EspC~EccA_1_ complex.

## CONCLUSION

In summary, we have purified and functionally characterized the EspC protein. The EspC enters and accumulated in the cytosol of human lung carcinoma A549 cells and caused the cell death. The purified EccA_1_ exits as a hexamer and interacts extensively with EspC export arm though its β-hairpin insertion of TPR domain. It is the first detailed characterization on biological activities and mechanism of recognition between of EspC and EccA_1_ proteins. Current knowledge will contribute significantly in the development of novel inhibitors, which will prevent the virulence proteins secretion by mycobacterial ESX-1 secretion system.

## AUTHOR INFORMATION

### Corresponding author

. Phone-011-26704155,

## Acknowledgements

Current project is supported by research grant from the Department of Science and Technology, India (No.-SR/SO/BB-0104/2012).Vipin K. Kashyap is supported by Senior Research Fellowship from UGC, India. Ruby Sharma is supported by Senior Research Fellowship from CSIR. Research grants from UGC-Networking and DST-PURSE from Jawaharlal Nehru University (JNU), New Delhi are gratefully acknowledged. The authors thank the staff members of Advanced Instrumentation Research Facility (AIRF), Jawaharlal Nehru University for their help in conducting the CD experiments.

## Author contributions

Vipin K. Kashyap was involved in purification of the EspC and EccA_1_ proteins, performed the EspC cell lysis and EccA_1_ ATPase assay, and performed binding analysis with native and mutant EspC and EccA_1_ proteins. Ruby Sharma prepared all EspC point mutants, purified all proteins and performed binding analysis with native and mutant EspC and EccA_1_ proteins. Ruby Sharma was involved in A549 cytotoxicity experiment and collected CD data on all EspC and EccA_1_ mutant proteins. During manuscript revision, Manoj and Abhishek were involved in A549 cellular cytotoxicity experiments. Ajay K. Saxena analyzed the entire data of all experiments, performed, CD data analysis, modeling and dynamics simulation of EspC-EccA_1_ complex, prepared all figures and wrote the entire manuscript.

## Conflict of interest

The authors declare they have no conflict of interest.

## ABBREVIATIONS

ATP: Adenosine Triphosphate
EspC: ESX-1 secreted protein C
EspB: ESX-1 secreted protein B
EccA1: ESX-1 conserved AAA+ATPase
CFP10: Culture Filtrate protein-10
ESAT6: Early Secretory Antigen Target-6
CD: Circular Dichroism
TPR: Tertratricopeptide repeat
OD: Optical Density
Ni-NTA: Nickel-Nitriloacetic Acid
SDS PAGE: Sodium Dodecyl Sulfate Polyacrylamide Gel Electrophoresis
SPR: Surface Plasmon Resonance
PBS buffer: Phosphate buffered saline
PBST buffer: Phosphate buffered saline with Tween-20.

## ACCESSION IDS

EspC protein UniProt ID: P9WJD7

EccA_1_ protein UniProt ID: P9WPH9

## Supplementary figures legends

**Figure S1**. The mass spectrometric analysis of purified EspC protein obtained from gel filtration column. The major peak corresponds to molecular weight of the EspC monomer ~11.9 kD.

**Figure S2. A,** Schematic diagram of various domains of EccA_1_ ATPase and its expression construct used for protein expression, TPR-tetratricopeptide repeat, ATPase-ATP binding domain. **B,** Size exclusion chromatography and SDS-PAGE analysis on purified EccA_1_. The EccA_1_ eluted as hexamer from Superdex 200(16/60) column.

**Figure S3. A,** The EccA_1_ was incubated with [γ^32^P] ATP and ATPase activity was determined at regular time intervals. The amount of hydrolyzed ATP is shown as a percentage of original [γ^32^P] ATP. Each point is the average of three independent experiments. The inset shows the plot of ATP hydrolysis~protein concentration and autoradiography profile was obtained from thin layer chromatography. **B,** Michaelis-Menten plot of ATP hydrolysis of the EccA_1_.

**Figure S4.** Molecular model of EspC hexamer was obtained by C_n_ symmetry module of ROSETTADOCK protein-protein docking server using EspC dimer as input model. The lowest energy cluster was chosen as best model and export arms of EspC hexamer are shown as the circle.

## Notes

### Competing Interest Statement

The authors have declared no competing interest.

