## Supplementary figures and images for "Dissection of function and recognition mechanism of *M. tuberculosis* ESX-1 secreted virulence factor EspC"

### Fig.S1

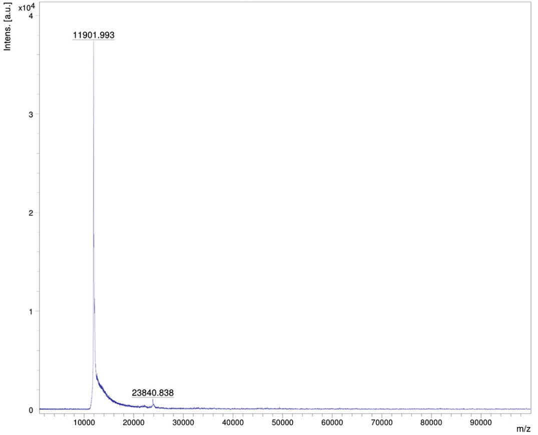

### Fig.S2

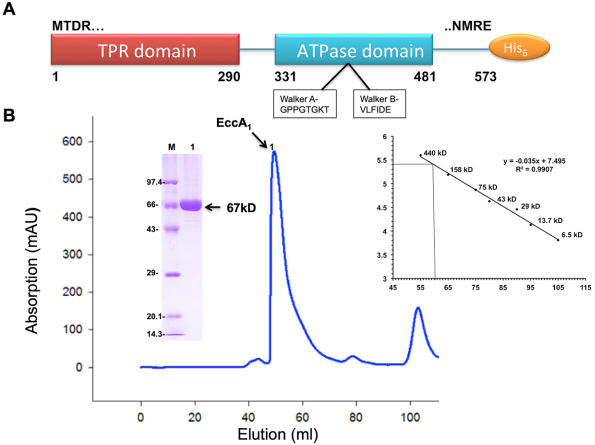

### Fig.S3

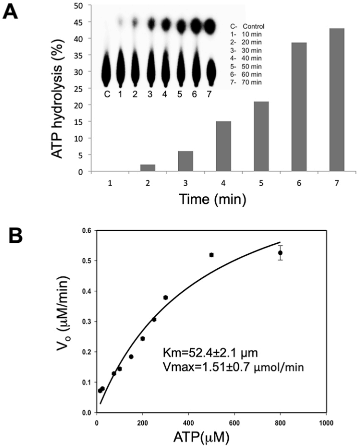

### Fig.S4

Fig. S4

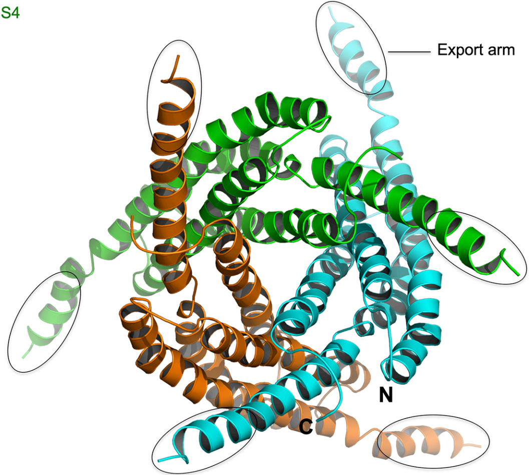
